# Loss of tau and Fyn reduces compensatory effects of MAP2 for tau and reveals a Fyn-independent effect of tau on glutamate-induced Ca^2+^ response

**DOI:** 10.1101/567776

**Authors:** Guanghao Liu, Ramasamy Thangavel, Jacob Rysted, Yohan Kim, Meghan B Francis, Eric Adams, Zhihong Lin, Rebecca J Taugher, John A Wemmie, Yuriy M Usachev, Gloria Lee

**Affiliations:** Interdisciplinary Program in Neuroscience, University of Iowa Carver College of Medicine, Iowa City, IA 52242, USA; Department of Internal Medicine, University of Iowa Carver College of Medicine, Iowa City, IA 52242, USA; Department of Pharmacology, University of Iowa Carver College of Medicine, Iowa City, IA 52242, USA; Department of Psychiatry, University of Iowa Carver College of Medicine, Iowa City, IA 52242, USA

**Keywords:** tau, Fyn, MAP2, calcium, proximity ligation assay

## Abstract

Microtubule-associated protein tau associates with Src family tyrosine kinase Fyn. A tau-Fyn double knockout (DKO) mouse was generated to investigate the role of the complex. DKO mice resembled Fyn KO in cognitive tasks and resembled tau KO mice in motor tasks and protection from pentylenetetrazole-induced seizures. In Ca^2+^ response, Fyn KO was decreased relative to WT and DKO had a greater reduction relative to Fyn KO, suggesting that tau may have a Fyn-independent role. Since tau KO resembled WT in its Ca^2+^ response, we investigated whether MAP2 served to compensate for tau, since its level was increased in tau KO but decreased in DKO mice. We found that like tau, MAP2 increased Fyn activity. Moreover, tau KO neurons had increased density of dendritic MAP2-Fyn complexes relative to WT neurons. Therefore, we hypothesize that in the tau KO, the absence of tau would be compensated by MAP2, especially in the dendrites, where tau-Fyn complexes are of critical importance. In the DKO, decreased levels of MAP2 made compensation more difficult, thus revealing the effect of tau in the Ca^2+^ response.

**Summary Statement:** The downstream effect of the interaction between microtubule-associated protein tau and Src family non-receptor tyrosine kinase Fyn was investigated with a tau/Fyn double KO mouse. We demonstrate that tau has a Fyn-independent role in glutamate-induced calcium response and that MAP2 can compensate for tau in interacting with Fyn in dendrites.

## Introduction

Tau was initially discovered as a microtubule-associated protein enriched in axons (Binder et al., 1986; Weingarten et al., 1975). Interest in tau dramatically increased after hyperphosphorylated tau was found in neurofibrillary tangles (NFT), a major hallmark of Alzheimer’s disease (AD) (Grundke-Iqbal et al., 1986; Kosik et al., 1986; Nukina and Ihara, 1986; Wood et al., 1986), and after mutations in the tau gene were found to cause frontotemporal dementia with Parkinsonism linked to chromosome 17 (Clark et al., 1998; Hutton et al., 1998; Poorkaj et al., 1998; Spillantini et al., 1998). Subsequent studies have revealed new functions for tau, such as the regulation of motor-driven axonal transport (Dixit et al., 2008; Trinczek et al., 1999), formation and trafficking of stress granules (Vanderweyde et al., 2016), and postsynaptic scaffolding (Ittner et al., 2010).

However, despite the multiple roles played by tau, initial studies of five lines of independently generated tau KO mice did not suggest any gross deficits or loss of viability(Dawson et al., 2001; Fujio et al., 2007; Harada et al., 1994; Tan et al., 2018; Tucker et al., 2001). Further testing of tau KO mice has yielded controversial results, as some studies reported minor memory deficits(Ahmed et al., 2014; Ikegami et al., 2000; Lei et al., 2014; Ma et al., 2014) while a larger number of studies reported no cognitive deficits(Dawson et al., 2010; Dawson et al., 2001; Ittner et al., 2010; Kimura et al., 2014; Lei et al., 2012; Li et al., 2014; Morris et al., 2013; Regan et al., 2015; Roberson et al., 2007; Tan et al., 2018; van Hummel et al., 2016). Morphological studies of tau KO mice reported alterations in small caliber axons as the only abnormality(Harada et al., 1994) while low density neuronal cultures from another line of tau KO mouse indicated slowed axonal development, although brain development appeared normal(Dawson et al., 2001). The lack of more pronounced behavioral and morphological phenotypes has largely been attributed to functional compensation by other microtubule-binding proteins such as microtubule associated protein 1A/1B (MAP1A/1B) and MAP2(Harada et al., 1994; Ma et al., 2014). While it is thought that compensating for a loss of microtubule stabilizing activity is important, given the additional functions for tau, other compensatory mechanisms may be needed if these functions are critical for mouse viability and depend on tau.

Following our finding that tau could associate with the cytoplasmic face of the plasma membrane(Brandt et al., 1995), we found that the proline-rich domain of tau interacts with the SH3 domain of Src family non-receptor tyrosine kinases (SFK) such as Fyn and Src(Lee et al., 1998) and that Fyn phosphorylated tau on Tyr18(Lee et al., 2004). In addition, we showed that tau increased the auto-phosphorylation of Fyn as well as the enzymatic activity of Fyn(Sharma et al., 2007). Also, tau was critical for nerve growth factor (NGF)-induced mitogen activated protein kinase (MAPK) activation, suggesting a role for tau in signal transduction (Leugers et al., 2013; Leugers and Lee, 2010). Interest in the tau-Fyn interaction has been increased by the presence of phospho-tyr18-tau in NFTs(Bhaskar et al., 2010; Lee et al., 2004), reported alterations of Fyn expression in AD brain(Ho et al., 2005; Shirazi and Wood, 1993), and the neuroprotective effects of Fyn depletion in AD models(Chin et al., 2004; Lambert et al., 1998). Furthermore, Fyn is highly expressed in the central nervous system(Umemori et al., 1992) and phosphorylates the N-methyl-D-aspartate receptor (NR) subunit 2B (NR2B) at Y1472 residue(Nakazawa et al., 2001; Rong et al., 2001), which facilitates the interaction between the NR and the postsynaptic density 95 protein (PSD-95)(Tezuka et al., 1999) and regulates long-term potentiation (LTP) and memory formation(Kojima et al., 1997; Stein et al., 1992). Tau has been implicated in targeting Fyn to the postsynaptic space, as tau KO mice showed decreased levels of Fyn and pY1472 NR2B in synaptosome preparations(Ittner et al., 2010). As a result, the PSD-NR interaction was disrupted, leading to protection from both pentylenetetrazole (PTZ) induced seizures and amyloid-β induced excitotoxicity (Ittner et al., 2010; Roberson et al., 2007). However, electrophysiological studies of tau KO resembled WT (Ittner et al., 2010; Kimura et al., 2014; Roberson et al., 2011), further underlying the need to elucidate the function of the tau-Fyn interaction and its significance in neuronal cell function.

To further understand the impact of the combined action of tau and Fyn, we generated a tau-Fyn double knockout (tau^−/−^/Fyn^−/−^) mouse, characterized its behavioral, biochemical, and neurophysiological properties, and found a role for tau in the neuron’s calcium response in a Fyn-independent manner. Additionally, we identified the location of tau-Fyn complexes in neurons and determined that in dendrites, the loss of such complexes led to compensation mediated by MAP2.

## Results

### Generation of tau-Fyn DKO mice

In order to investigate the downstream effects of the tau-Fyn interaction, we generated tau^−/−^/Fyn^−/−^ double knockout mice (DKO) by breeding tau KO (C57BL/6) mice(Dawson et al., 2001) with Fyn KO (C57BL/6-S129) mice(Stein et al., 1992) (Fig. S1). Polymerase chain reaction (Fig. 1A) was used for genotyping and western blotting (Fig. 1B) was used to confirm the loss of tau and Fyn.

**Fig. 1:**
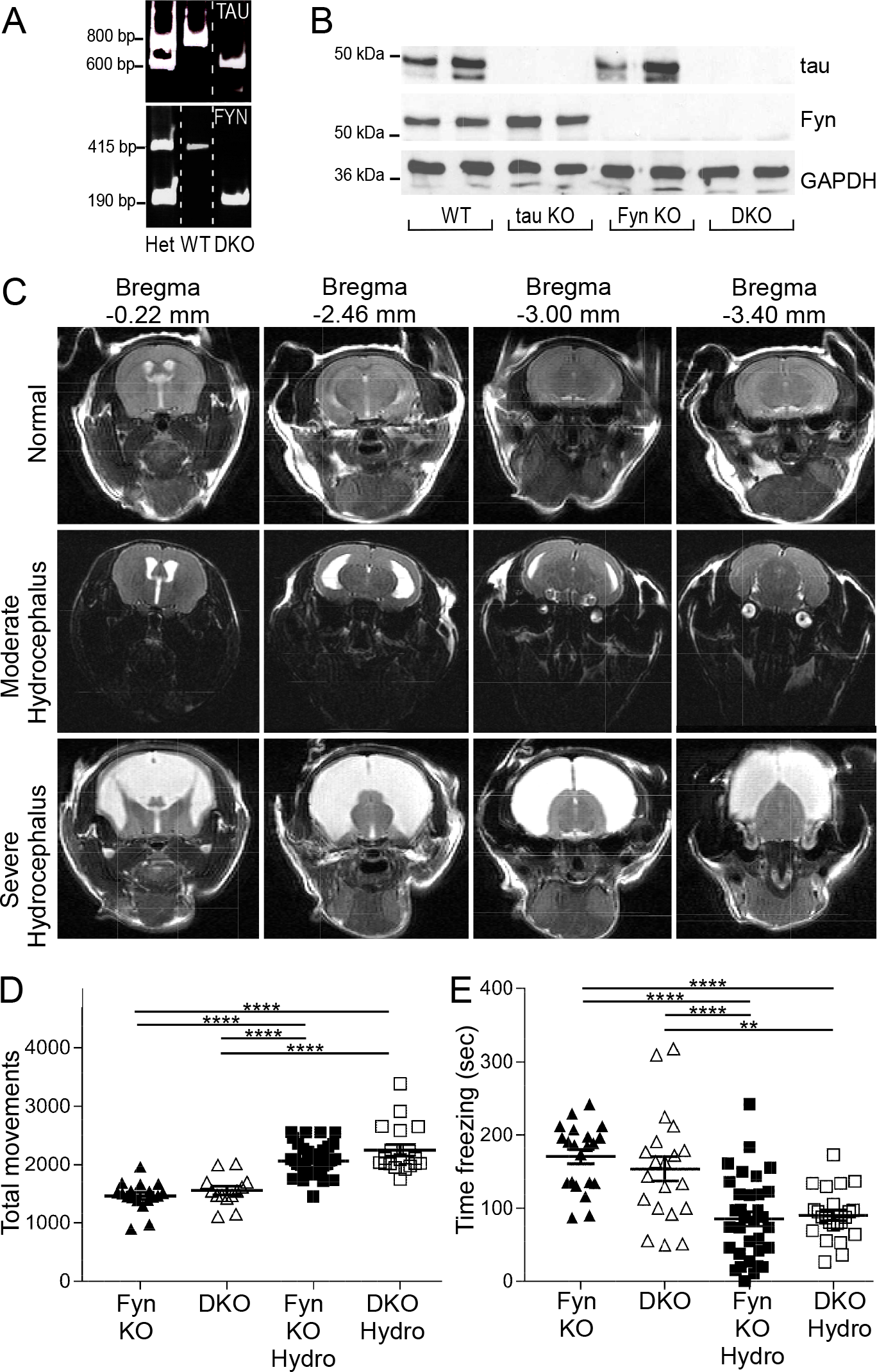
Confirmed by MRI, tau^−/−^/Fyn^−/−^ double knockout (DKO) mice with hydrocephalus exhibited behavioral abnormalities. A) Polymerase chain reaction (PCR) products from heterozygous (Het), WT, and DKO mice are shown. B) Western blot of crude brain lysate showed that DKO mice did not express tau or Fyn. C) Examples of T2 weighted MRI of coronal mouse brains showing normal, moderate, or severe hydrocephalus. D) In the open field test, Fyn KO and DKO mice with hydrocephalus (moderate or severe) had increased total movements relative to mice with no hydrocephalus. n=18 Fyn KO, 13 DKO, 41 Fyn KO hydro, 19 DKO hydro. ****p≤ 0.0001. E) In contextual fear conditioning, Fyn KO and DKO mice with moderate or severe hydrocephalus had decreased freezing relative to mice with no hydrocephalus. n=21 Fyn KO, 20 DKO, 33 Fyn KO hydro, 23 DKO hydro. **p=0.0014, ****p≤ 0.0001. Mean ± S.E.M. are shown. Ordinary one-way ANOVA with Tukey’s post-hoc multiple comparison was used for both panels D and E.

During breeding, DKO animals exhibited a propensity to develop non-obstructive hydrocephalus postpartum, consistent with the observation that having the homozygous Fyn KO trait on a C57BL/6 background increased the occurrence of hydrocephalus (Goto et al., 2008). While mice with severe hydrocephalus could be identified within 6 weeks of age by their domed heads and hunched backs, less severe levels of hydrocephalus escaped visual detection. Therefore, to identify mice with moderate hydrocephalus, MRI was used to image the lateral ventricle. Quantification of the images allowed us to classify mice as being normal or having moderate or severe hydrocephalus (Fig. 1C). Behavioral tests showed that both Fyn KO and DKO mice with mild and severe hydrocephalus had increased total movements in the open field test (Fig. 1D) and decreased total freezing in contextual fear conditioning (Fig. 1E) when compared to normal mice of the same genotype. This suggested that hydrocephalus caused the animal to be hyperactive. Thus, subsequent behavioral and biochemical tests were performed using only non-hydrocephalic mice with normal brain structure that had been screened by MRI. Beginning at eight weeks of age, the mice underwent a series of motor and behavioral tests.

### DKO mice recapitulate specific behaviors of Fyn KO and tau KO mice

Using the open field apparatus, mice of the four genotypes had similar total movements and percent movement in the center (Fig. S2A, B). This indicated an absence of motor deficits or anxiety phenotype at young age. However, when mice were aged to 12 months and underwent the pole test, tau KO and DKO mice required more time to descend the pole relative to WT and Fyn KO mice (Fig. S2C).

In novel object recognition, mice from the four genotypes displayed equal preferences for two identical objects on training day (Fig. 2A, left). However, on testing day, both Fyn KO and DKO mice performed significantly worse than WT and tau KO mice in interacting with the new object (Fig. 2A, right). There were no differences between WT and tau KO mice (p=0.9012), and between Fyn KO and DKO mice (p=0.1301). The absence of anxiety as monitored by the open field test suggested that Fyn KO and DKO had a cognitive deficit.

**Fig. 2:**
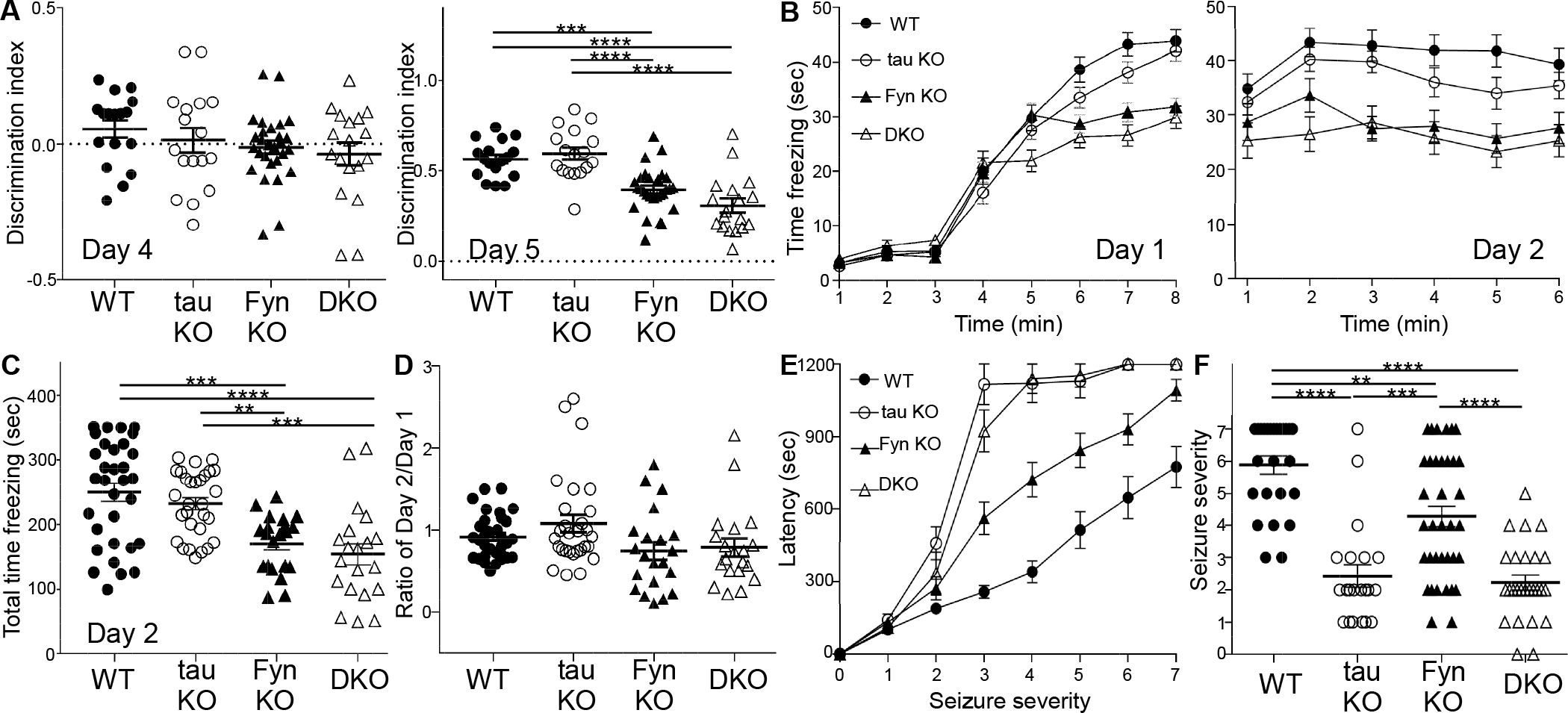
In memory tasks, DKO recapitulated the behavior of Fyn KO mice whereas in PTZ induced seizures, DKO mimicked the tau KO mice. A) In novel object recognition, there was no difference in object preference on training day (One-way ANOVA p= 0.3221; left panel) while on testing day, Fyn KO and DKO mice had significantly reduced interaction with the novel object (right panel; One-way ANOVA: p<0.0001; ***p=0.0004, ****p≤ 0.0001). n= 17 WT, 17 tau KO, 28 Fyn KO, 18 DKO. B) In contextual fear conditioning, using minute by minute measurements, WT and tau KO mice spent more time freezing compared to Fyn KO and DKO mice on both training day (left) and testing day (right). n= 32 WT, 29 tau KO, 21 Fyn KO, 20 DKO. C) Fyn KO and DKO mice spent less time freezing than WT (p=0.0002, p<0.0001, respectively) and tau KO mice (p=0.0075, p=0.0004, respectively). One-way ANOVA: p<0.0001; **p= 0.0075, ***p= 0.0002 or 0.0004, ****p≤ 0.0001. D) Ratio of time freezing in first minute of day 2 to time freezing in last minute of day 1 show an average of 1 for all genotypes, indicating a learning rather than memory deficit for Fyn KO and DKO mice. One-way ANOVA: p=0.0509 E) In PTZ induced seizures, tau KO and DKO mice were equally protected while Fyn KO mice were only moderately protected as measured in latency to reach each seizure stage. n= 25 WT, 21 tau KO, 36 Fyn KO, and 26 DKO. F) For maximum seizure stage reached, tau KO and DKO mice were similarly protected relative to WT and Fyn KO mice; Fyn KO mice were only slightly protected relative to WT mice. One-way ANOVA: p<0.0001; **p= 0.0011, ***p= 0.0003, ****p≤ 0.0001. n= 25 WT, 21 tau KO, 36 Fyn KO, and 26 DKO. Mean ± S.E.M. are shown. Ordinary one-way ANOVA with Tukey’s post-hoc multiple comparisons were used for A, B, C, D, F. Ordinary two-way ANOVA with Tukey’s post-hoc multiple comparisons were used for B and E.

In contextual fear conditioning, Fyn KO and DKO mice spent significantly less time freezing as compared to WT and tau KO mice on training day (Fig. 2B, left). On testing day, Fyn KO and DKO mice also spent less time freezing than WT and tau KO mice as measured using either minute by minute measurements (Fig. 2B right panel) or total time freezing (Fig. 2C). There was no statistically significant difference between Fyn KO and DKO mice (p=0.8442) and between WT and tau KO mice (p=0.7076) (Fig. 2C). To assess learning versus memory, the amount of freezing during the first minute of the testing day was divided by the amount of freezing during the last minute of the training day. There was no significant difference between the ratios of the four genotypes (Fig. 2D), indicating that the mice did not differ with respect to memory and that the deficits exhibited by Fyn KO and DKO mice on day 2 were due to a learning deficit.

Previous reports have also shown that tau ablation protected against pentylenetetrazole (PTZ) induced seizures (Roberson et al., 2007). This property has been attributed to tau depletion reducing levels of Fyn at the post-synaptic region leading to disrupted NR stability within the post-synaptic density(Ittner et al., 2010). In terms of both latency to develop seizures and maximum seizure stages reached, DKO and tau KO mice were equally and dramatically protected against PTZ whereas Fyn KO mice were only moderately protected relative to WT mice (Fig. 2E, F). There was no significant difference between tau KO and DKO mice (p=0.9752; Fig. 2F). To summarize, in motor tasks and PTZ induced seizures, DKO mice resembled tau KO mice while in cognitive and memory tasks, DKO mice resembled Fyn KO mice.

### Localization of Tau-SFK complexes in WT primary hippocampal culture

To further investigate the functions of the tau/Fyn complex, proximity ligation assays (PLA) were used to localize tau-Fyn complexes in WT primary hippocampal cultures (Fig. 3A). PLAs are able to detect endogenous protein-protein interactions *in situ;* signals are recovered when the two PLA probes lie within 40 nm of each other, using rolling circle amplification to yield a marker for the interaction (Gullberg et al., 2004) (Soderberg et al., 2006). As a control, tau KO cultures were similarly probed (Fig. S3B). Tau-Fyn complexes were found in all cellular compartments of WT hippocampal neurons (axons (Fig. 3B), dendrites (Fig. 3C), and cell bodies (Fig. 3D)). For quantitation, PLA dots were counted in both dendrites and axons. Dot intensity was not used because the intensity of a PLA signal reflected the rolling circle step, making the intensity level an artifical characteristic. Upon quantitation, the density of tau-Fyn complexes in dendrites (number of PLA puncta/process length) was 2.54 fold higher than that in axons (Fig. 3E). As expected, there were no tau-Fyn complexes in tau KO neurons (Fig. S3A). Since we have previously shown that tau increased Fyn auto-phosphorylation(Sharma et al., 2007), complexes between tau and activated SFK (pSFK) were also examined (Fig. 3F-J). Similar to the tau-Fyn complexes, the density of tau-pSFK complexes in dendrites was 2.04 fold higher than that in axons (Fig. 3J). However, since the pSFK antibody cannot distinguish between activated Fyn and activated Src, tau-Src complexes were also examined as a control. The density of tau-Src complexes in dendrites was increased by 2.78 fold relative to that in the axon (Fig. 3O).

**Fig. 3:**
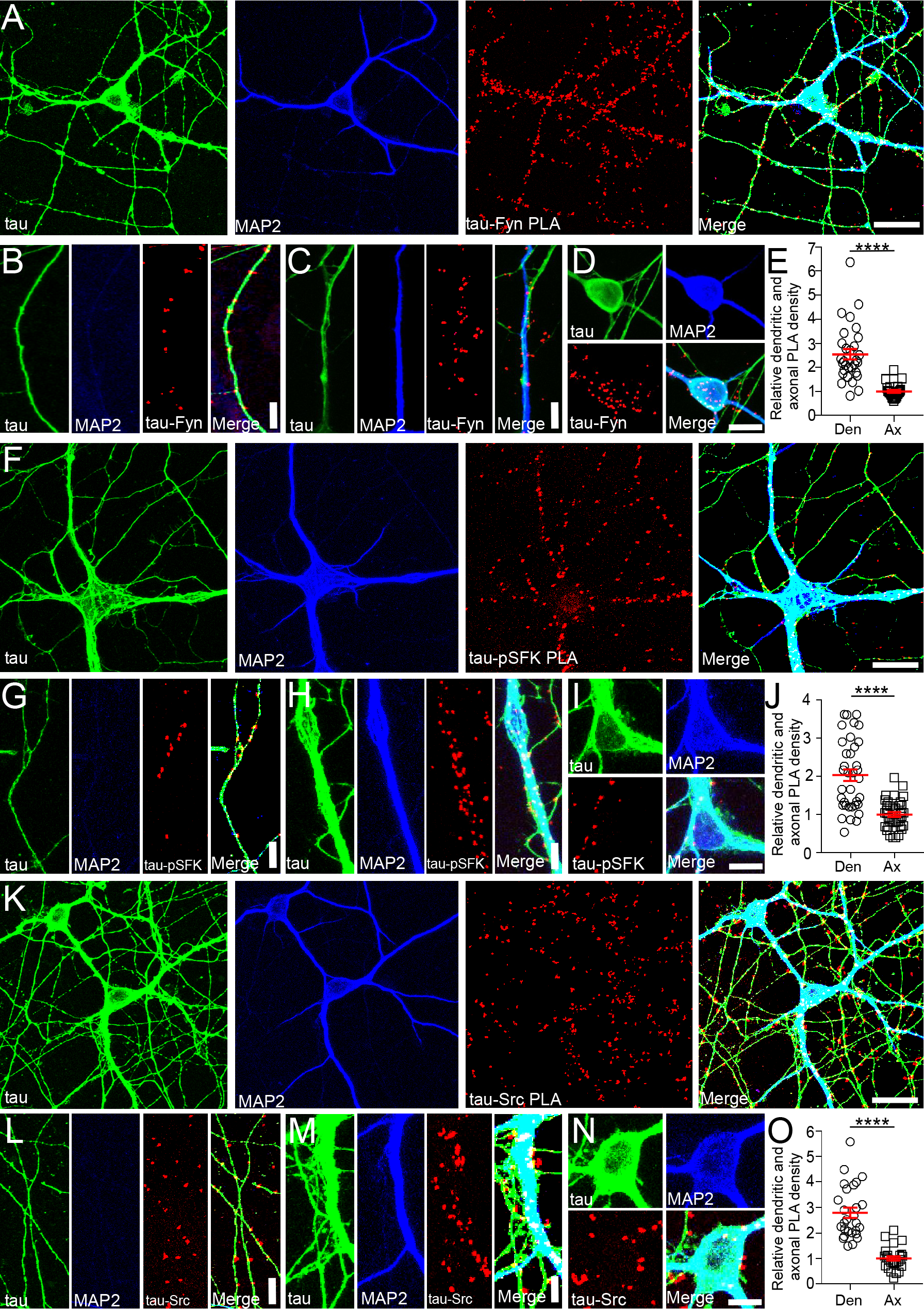
Tau-Fyn, tau-pSFK, and tau-Src complexes are present in neurons. A,F,K) Using WT hippocampal neurons, tau-Fyn (A), tau-pSFK (F), and tau-Src (K) complexes were identified using proximity ligation assay as described in Materials and Methods. Scale bar: 25µm. B-D, G-I, L-N) Tau-Fyn, tau-pSFK, and tau-Src complexes were found in axons (B, G, L), dendrites (C, H, M), and cell bodies (D, I, N), respectively. Scale bar: 10µm (B, C, G, H, L, M); 20 µm (D, I, N). E, J, O) PLA complex density in WT neurons was quantitated as described in Materials and Methods. Mean ± S.E.M. are shown. n=30 areas from each condition were analyzed. ****p≤ 0.0001 as determined by unpaired parametric 2 tailed t-test.

Since tau is known to be enriched in axons, we found it surprising that the density of tau-SFK complexes was higher in the dendrites relative to axons. However, we noted that axons were significantly longer and more numerous than dendrites and that the density of PLA puncta was relatively high in the proximal axon but dramatically decreased in the distal axon whereas the density of PLA puncta in the shorter dendrites was uniformly high. In fact, if only the proximal axon was used to calculate density, the density of tau-Fyn complexes in axons would match that of the dendrites (Fig. S3). Therefore, when the total axon length was used, the calculated density in axons was lower than that in dendrites.

### Fyn KO and DKO hippocampal insoluble PSD fractions have decreased phospho-Y1472 NR2B and phospho-SFK levels

Because the density of tau-Fyn complexes in dendrites exceeded that in axons and because tau has been shown to target Fyn to the post-synaptic density to affect NR2B phosphorylation (Ittner et al., 2010), we wanted to determine if NR2B phosphorylation was altered in the DKO mice. Crude hippocampal synaptosomes were fractionated into “soluble non-PSD” and “insoluble PSD” fractions using Triton X-100 (Fig. 4A) (Ittner et al., 2010; Lopes et al., 2016b; Milnerwood et al., 2010). The presence of synaptophysin in the “soluble non-PSD” and the presence of PSD-95 in the “insoluble PSD” fractions (Fig. 4B) demonstrated the separation of PSD and non-PSD membranes from crude WT synaptosomes; tau and Fyn were present in both fractions (Fig. 4B). In examining the “insoluble PSD” fraction from 9-12 month old hippocampus, PSD-95 and total NR2B levels were not significantly different between the four genotypes (One-way ANOVA p=0.6462, p=0.3439, respectively; Fig. 4C, D). However, relative to WT, pY1472-NR2B levels were decreased by 54.6 % in Fyn KO and 64.0 % in DKO insoluble PSD fractions, with Fyn KO and DKO not being significantly different from each other (p=0.8871; Fig. 4C, D). WT and tau KO were also not significantly different from each other (p=0.8284; Fig. 4C, D). We then examined whether Src might be compensating for Fyn. In the PSD fraction, although the Src level was unchanged in the four genotypes (p=0.7699; Fig. 4C, D), the activated SFK (pSFK) level in Fyn KO and DKO was decreased by 73.2% and 84.0% with respect to that of WT. There was no significant difference between tau KO and WT (p=0.9946) and between Fyn KO and DKO (p=8512; Fig. 4C, D). While the pSFK antibody cannot distinguish between activated Fyn and activated Src, all four genotypes had similar Src levels in the PSD but only Fyn KO and DKO had decreased pSFK levels. Since the absence of Fyn did not alter Src levels but reduced pSFK levels, we concluded that most of the activated tyrosine kinase in WT and tau KO PSD fractions was Fyn that was responsible for phosphorylating Y1472-NR2B.

**Fig. 4:**
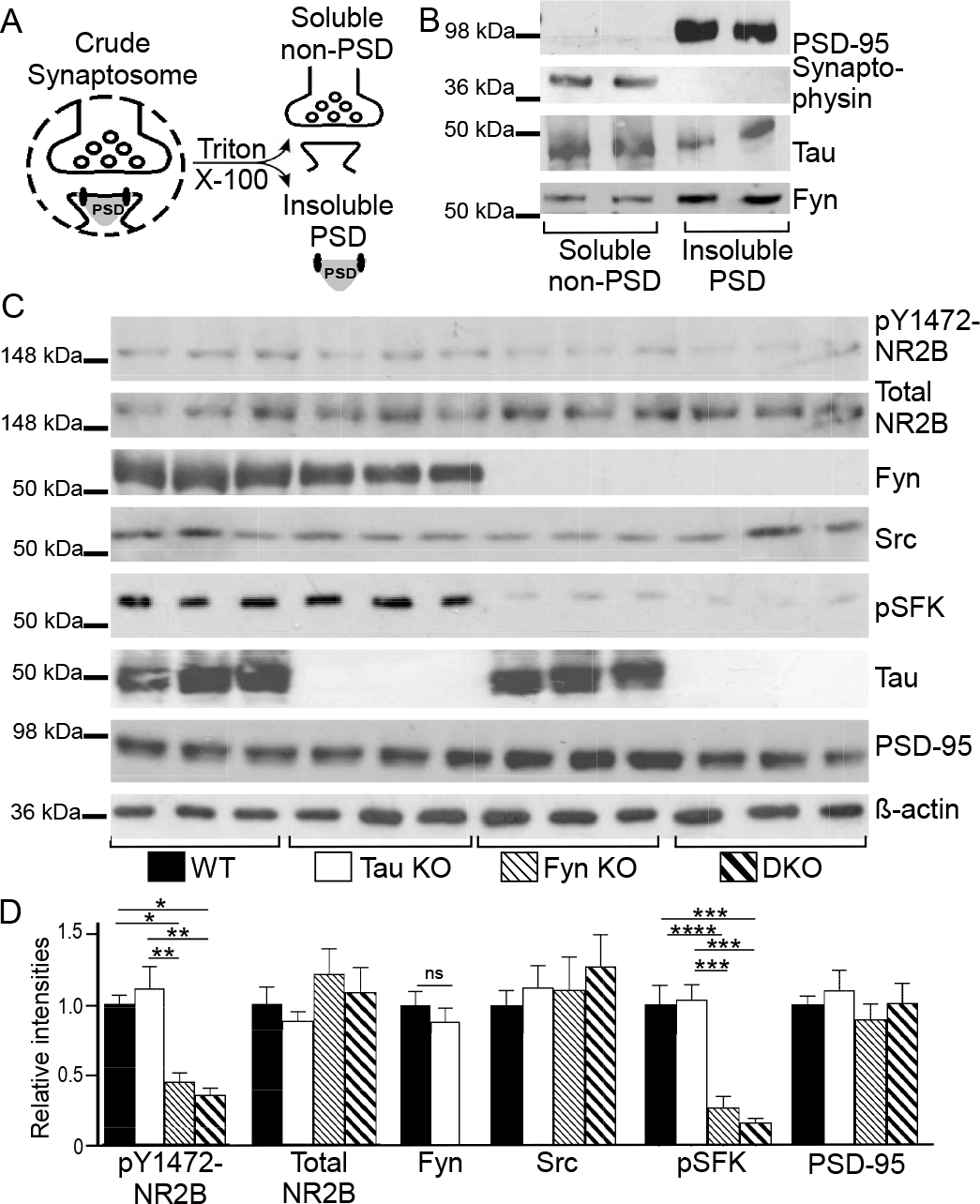
Fyn KO and DKO hippocampal PSD fraction had decreased phospho-Y1472 NR2B and phospho-SFK levels. A) Schematic for separating insoluble PSD and soluble non-PSD fractions from crude synaptosomes. B) Soluble non-PSD fraction isolated from crude hippocampal synaptosomes of WT mice showed synaptophysin, a pre-synaptic marker, while insoluble PSD fraction showed PSD95, a post-synaptic marker. Fyn and tau were found in both fractions. C) Western blotting of hippocampal insoluble PSD fractions prepared from WT, tau KO, Fyn KO, and DKO. D) Blots were quantitated by densitometry and normalized. n=9 animals for each genotype. For pY1472-NR2B, relative to WT, levels were decreased in Fyn KO and in DKO (***p=0.0008, ****p<0.0001, respectively); relative to tau KO, levels were also significantly decreased in Fyn KO and in DKO (****p0.0001, ****p<0.0001, respectively). For pSFK, relative to WT, levels were decreased in Fyn KO and in DKO (****p<0.0001, ****p<0.0001, respectively); relative to tau KO, levels were also decreased in Fyn KO and in DKO (****p<0.0004, ****p<0.0004, respectively). Mean ± S.E.M. are shown. Unpaired two-tailed parametric t-test was used for Fyn and ordinary one-way ANOVA with Tukey’s post-hoc multiple comparisons was used for others.

### Decreased glutamate-induced Ca^2+^ response in Fyn KO and DKO primary hippocampal neurons

Since phosphorylation of NR2B has been shown to be involved in facilitating Ca^2+^ response upon NR activation(Nakazawa et al., 2001; Rong et al., 2001; Tezuka et al., 1999), we were interested in determining the glutamate-induced Ca^2+^ influx of DKO neurons. In addition, depleting tau with shRNA in WT neurons had already been shown to affect glutamate-induced, NMDA receptor-dependent Ca^2+^ influx (Miyamoto et al., 2017), so we were also interested in the response of tau KO neurons. Sample tracings of [Ca^2+^]_i_ from five individual cells of each genotype are shown in Fig. 5A. At baseline, unstimulated intracellular [Ca^2+^]_i_ from the four genotypes were similar (p=0.2808; Fig. 5B). Upon stimulation, Fyn KO neurons had a 29.9% reduction and DKO neurons had a 53.1% reduction relative to WT neurons; tau KO neurons were not significantly different from WT neurons (p=0.7264). However, Ca^2+^ response in DKO neurons was significantly different from Fyn KO neurons (Fig. 5C).

**Fig. 5:**
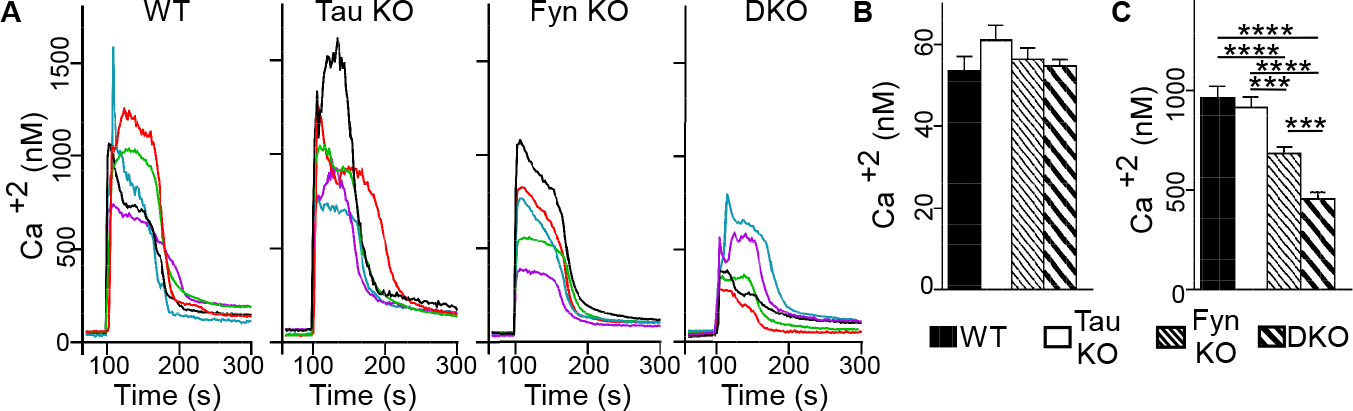
Glutamate induced Ca^2+^ response was impaired in Fyn KO and DKO hippocampal neurons. A) Ca^2+^ response traces from five representative neurons from each genotype are shown; traces were obtained in response to glutamate and glycine as described in Materials and Methods. B) No difference in baseline intracellular [Ca^2+^]_i_ was found among the hippocampal neurons from the different genotypes. n= 105 WT, 115 tau KO, 114 Fyn KO, and 181 DKO cells were analyzed. p=0.2808. **C)** Upon stimulation, Fyn KO neurons had a 29.9% reduction (****p<0.0001) and DKO neurons had a 53.1% reduction (****p<0.0001) in Ca^2+^ response relative to WT neurons. The reduction of the DKO neurons was significantly different from that of Fyn KO neurons (***p=0.0001). Mean ± S.E.M. are shown. Ordinary one-way ANOVA with Tukey’s post-hoc multiple comparisons was used.

The fact that DKO neurons had an even greater reduction relative to Fyn KO neurons suggests that tau had a Fyn-independent role in regulating Ca^2+^ response that was unmasked in the absence of Fyn. In addition, the absence of any changes in the Ca^2+^ response of tau KO neurons suggested that another protein could compensate for tau in the tau KO. However, this compensatory effect was apparently lost in the DKO, leading to an exacerbated decrease in Ca^2+^ influx.

### Microtubule-associated protein 2 (MAP2) was altered in KOs and like tau, could increase Fyn activity

Our finding that tau KO had no memory deficits, no changes in pY1472-NR2B, and no changes in glutamate-induced Ca^2+^ response suggested that either Fyn did not require tau to mediate these functions or that a different protein was compensating for the loss of tau with respect to its Fyn interaction. Interestingly, another microtubule-associated protein, MAP2, has been found to be upregulated in tau KO mice (Ma et al., 2014) and in agreement with the previous data, we also found MAP2 to be increased by 48.9% in the tau KO mice (p=0.021, Fig. 6A). However, while it has been shown that MAP2 compensated for the loss of tau by maintaining microtubule stability, as measured by tubulin acetylation (Ma et al., 2014), the possibility of other compensating mechanisms remained unexplored.

**Fig. 6:**
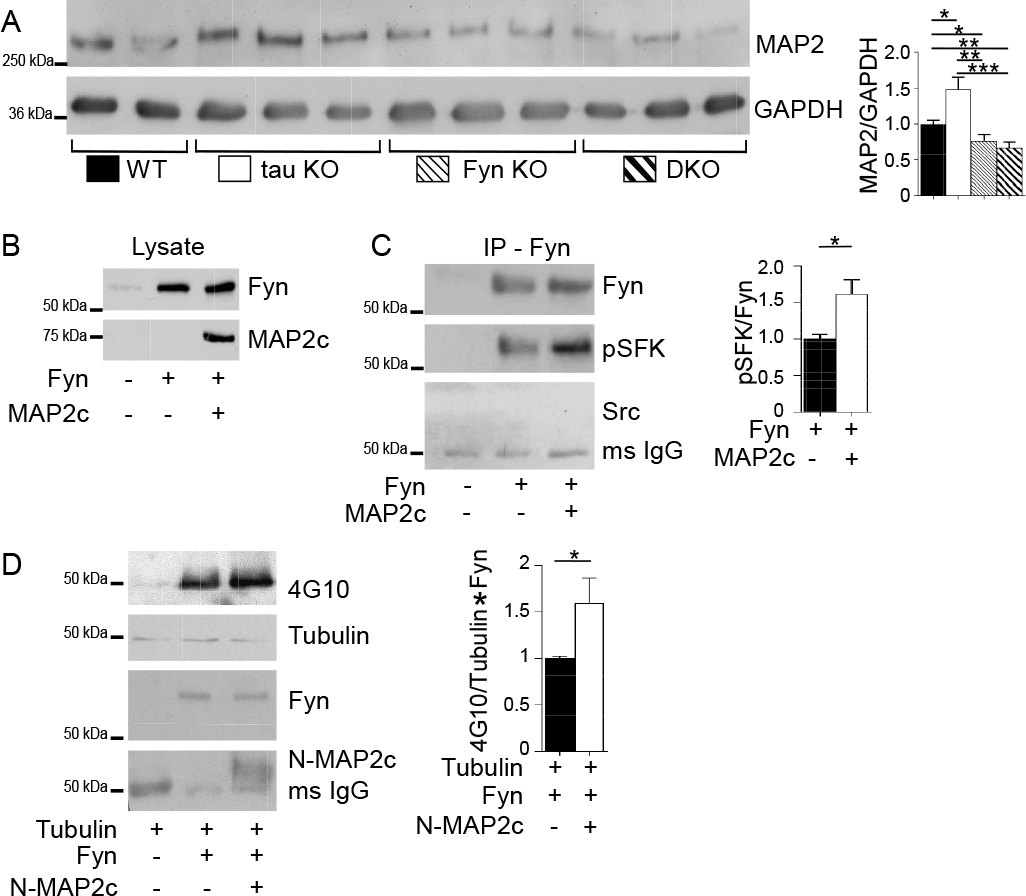
Microtubule-associated protein 2 (MAP2) is increased in tau KO and its association with Fyn increases Fyn activity. A) Crude mouse brain lysates from four genotypes were probed for MAP2, with GAPDH as control. Relative to WT mice, MAP2 was increased in tau KO mice (* p=0.021), decreased in Fyn KO mice (*p=0.044) as well as in DKO mice (**p=0.0045). Mean ± S.E.M. are shown 7 WT, 8 tau KO, 8 Fyn KO, and 8 DKO mice were used. B) Lysates from 3T3 cells confirmed transfection with Fyn alone or Fyn plus MAP2c. C)Fyn IP from cells co-transfected with Fyn plus MAP2c showed increased pSFK levels relative to Fyn IP from cells transfected with Fyn alone. No Src was detected (C). n=4 independent experiments were conducted. *p=0.0254. D) Tubulin was incubated either alone, with Fyn, or with Fyn plus N-MAP2c, then probed with anti-phospho-tyrosine (4G10). n=6 independent experiments. Unpaired two-tailed parametric t-test was used for A, C, D. Mean ± S.E.M. are shown.

Similar to tau, MAP2 has a PXXP domain that interacts with the SH3 domain of Fyn (Zamora-Leon et al., 2001). While we have shown that for tau, this interaction potentiated Fyn and Src activities (Sharma et al., 2007), the ability of MAP2 to exhibit a similar property has not been demonstrated. To investigate, MAP2c, the shortest isoform in the MAP2 family was tested for its ability to increase auto-phosphorylation of Fyn and to increase the enzymatic activity of Fyn. To examine auto-phosphorylation of Fyn, as detected by antibodies against activated SFK, 3T3 cells were transfected with Fyn alone or with MAP2c (Fig. 6B). After Fyn was immunoprecipitated (IPed) and probed for activated SFK, we found that the phosphorylation of Fyn expressed with MAP2c was increased by 61.9% relative to Fyn expressed alone (Fig. 6C). This indicated that the presence of MAP2c increased the auto-phosphorylation of Fyn. To ensure that the anti-Fyn antibody was specific for Fyn and not other SFKs, the IPed Fyn was also probed for Src. Src was not detected (Fig. 6C), indicating that the IP was indeed specific for Fyn. To investigate the ability of MAP2c to increase the enzymatic activity of Fyn, *in vitro* kinase assays were conducted using tubulin, a known substrate for Fyn (Marie-Cardine et al., 1995). Because MAP2c can bind tubulin, an N-terminal fragment of MAP2c (N-MAP2c) containing the Fyn binding motif but lacking the microtubule binding domain was used. Fyn and N-MAP2c were separately expressed in 3T3 cells and IP’ed for use in the kinase reaction. Tubulin was incubated either alone, with Fyn, or with pre-incubated Fyn and N-MAP2c. The presence of N-MAP2c increased tyrosine phosphorylation of tubulin by 59.4% relative to tubulin incubated with Fyn alone (p=0.049; Fig. 6D). Our data provide evidence that similar to tau, MAP2 can increase Fyn activity. As an additional indication that MAP2 resembled tau with regards to potentiating SFK activity, we examined the ability of MAP2c to sustain SFK activation in stimulated 3T3 cells. In this system, SFK activation was indirectly assessed and we had previously shown that tau was able to prolong SFK activation following stimulation. When MAP2 was compared to tau in this assay, we found that MAP2 resembled tau, prolonging SFK activation in cells (Fig. S5).

Interestingly, when MAP2 levels were assessed in Fyn KO and DKO mice, MAP2 was decreased by 23.6% and 33.2 %, respectively, relative to WT mice (p=0.044 and p=0.0045; Fig. 6A), with was no significant difference between the two (p=0.438; Fig. 6A). The labeling of MAP2 in 12-month-old hippocampal sections from the four genotypes confirmed the biochemical result (Fig. S4). Compared to WT, hippocampal sections of Fyn KO and DKO mice appeared to display shorter MAP2 fibers and weaker MAP2 intensity while tau KO mice appeared to show stronger MAP2 intensity (Fig. S4). Therefore, in regulating MAP2 expression levels, Fyn appeared to act downstream of tau, since the DKO resembled the Fyn KO. In the DKO, the decrease in MAP2 most likely eliminated the ability of MAP2 to compensate for the loss of tau. Thus, when DKO neurons experienced a further decrease in calcium influx, relative to the Fyn KO, we were able to identify a Fyn-independent effect that tau had on glutamate-induced calcium influx. In the tau KO, it was likely that this effect was not detected because of the increased MAP2 level.

### Tau KO neurons had more dendritic MAP2-Fyn complexes than WT neurons

To determine whether MAP2-Fyn complexes might compensate for the loss of tau-Fyn complexes in the tau KO, MAP2-Fyn and MAP2-pSFK complexes were visualized in WT and tau KO primary hippocampal cultures using PLA (Fig. 7). Relative to WT dendrites, tau KO dendrites had a 32.5% increase in MAP2-Fyn density (Fig. 7A, B, G) and a 39.9% increase in MAP2-pSFK density (Fig. 7C, D, H). As expected, we did not find any MAP2-containing complexes in axons, identified as MAP2 negative and tubulin positive processes (Fig. 7, open arrowheads). In tau KO neurons, MAP2-Src complex density was increased 36.8% relative to WT neurons (Fig. 7 E, F, I). Together, these data support the notion that MAP2 compensates for tau in terms of interacting with SFKs in dendrites.

**Fig. 7:**
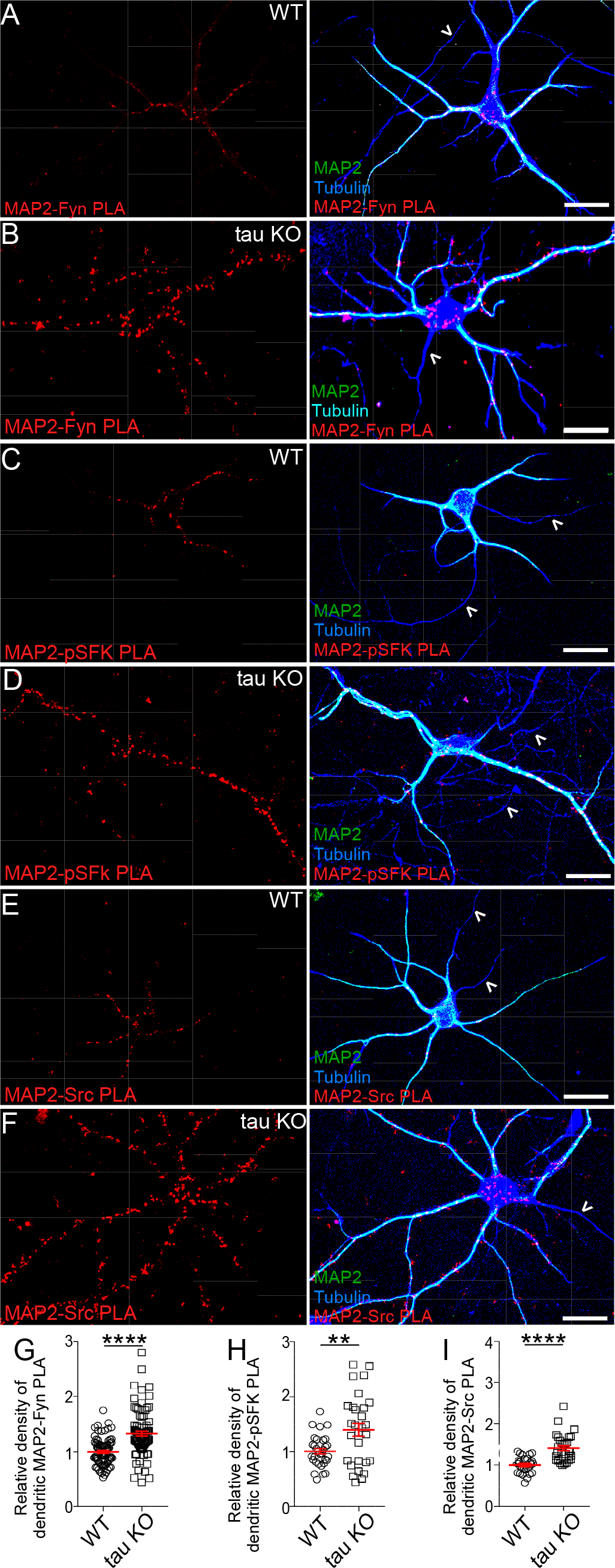
Tau KO neurons have more MAP2-Fyn, MAP2-pSFK, and MAP2-Src complexes relative to WT neurons. MAP2-Fyn (A, B), MAP2-pSFK (C, D), and MAP2-Src complexes (E, F) were detected in WT neurons (A,C,E) or tau KO neurons (B, D, F) using proximity ligation assay. No MAP2-SFK complexes were visualized in axons (open arrow heads). Scale bar: 25µm. MAP2-Fyn (G), MAP2-pSFK (H), and MAP2-Src (I) complex densities were quantitated in both WT and tau KO neurons. Mean ± S.E.M. are shown, 88 images for MAP2-Fyn, 30 images for MAP2-pSFK, and 30 images for MAP2-Src were analyzed. Unpaired parametric two-tailed t-test was used. For G) and I), ****p≤ 0.0001 and for H), **p=0.0033.

## Discussion

In the present study, we generated tau^−/−^/Fyn^−/−^ DKO mice to investigate the combined effect of tau and Fyn and found that they resembled tau KO mice in motor tasks and protection from PTZ while resembling Fyn KO mice in cognitive tasks. DKO and Fyn KO mice also had decreased Y1472-NR2B phosphorylation levels in hippocampal insoluble PSD fractions. These changes were accompanied by a reduction in glutamate-induced Ca^2+^ response in Fyn KO and DKO primary hippocampal neurons, with DKO neurons having a more severe reduction than Fyn KO neurons, indicating that tau may have a Fyn-independent effect on regulating Ca^2+^ influx. Together with our findings that MAP2 potentiated Fyn activity and that dendritic MAP2-Fyn complexes were increased in tau KO neurons relative to WT neurons, our data suggest that MAP2 may compensate for the loss of tau by increasing the levels of MAP2-Fyn and MAP2-pSFK complexes in dendrites.

In motor tasks, our aged tau KO and DKO mice had deficits in the pole test, which was consistent with results obtained by other laboratories (Ikegami et al., 2000; Lei et al., 2014; Lopes et al., 2016a; Morris et al., 2013). Since aged Fyn KO mice did not exhibit such deficits, we concluded that tau, not Fyn, was involved in the motor pathway. Another known property for tau KO mice is their protection from PTZ-induced seizures and Aβ induced excitotoxicity (Roberson et al., 2007), traits attributed to reduced Fyn levels at the post-synaptic region and disrupted NR stability within the post-synaptic density(Ittner et al., 2010). However, in PTZ-induced seizures, our finding that tau KO mice had greater protection than Fyn KO mice suggested that tau may participate in Fyn independent pathways to regulate seizure susceptibility. If tau acted only with Fyn, Fyn KO mice would have shown protection against PTZ equal to that of the tau KO. The fact that in response to PTZ, DKO and tau KO mice had similar seizure thresholds argued that Fyn’s protective effect against seizure was encompassed by tau’s protective effect and did not represent a separate pathway since the DKO mice did not exhibit an additional level of protection relative to tau KO mice. Besides interacting with Fyn and affecting NR2B-PSD95 association, tau has been reported to affect two other pathways that regulate seizure susceptibility. One pathway involves SynGAP1, a site-specific inhibitor of Ras, where it has been found that reducing SynGAP1 in tau KO mice increased susceptibility to PTZ-induced seizure (Bi et al., 2017). A second pathway has been reported where the loss of tau prevented the depletion of Kv4.2, a dendritic potassium channel, and mitigated hyperexcitability in an AD mouse model (Hall et al., 2015). These data indicate that tau is involved in other mechanisms that modulate seizure susceptibility and excitotoxicity in dendrites. We speculate that one or both of these mechanisms will be Fyn-independent.

Our data also provide evidence that Fyn plays a role in regulating PTZ seizure severity, as our non-hydrocephalic Fyn KO mice had decreased PTZ-induced seizure susceptibility. Our results disagreed with previous reports that Fyn KO mice had either increased seizure susceptibility(Miyakawa et al., 1996) or similar susceptibility(Kojima et al., 1998) relative to WT mice. However, these previous studies likely included mice with undetected mild/moderate hydrocephalus. By using MRI to identify Fyn KO mice with mild/moderate hydrocephalus, we had found that mice with hydrocephalus had increased seizure susceptibility compared to WT (Fig. S6). In addition, when a cohort of superficially normal Fyn KO mice, with and without hydrocephalus, was analyzed as a group, the seizure threshold resembled that of WT (Fig. S6). Therefore, because prior studies were not aware that the Fyn KO mice might have had mild/moderate hydrocephalus (Kojima et al., 1998; Miyakawa et al., 1996), their data did not reflect the sole contribution of the Fyn KO trait. Our data suggest that the Fyn KO trait conferred mild protection against seizures.

In agreement with prior studies where mice lacking Fyn, but not other members of Src family non-receptor tyrosine kinases, had deficits in LTP and performance in the Morris water maze (Grant et al., 1992), we found Fyn, not tau, to be critically involved in cognitive tasks(Dawson et al., 2010; Dawson et al., 2001; Ittner et al., 2010; Kimura et al., 2014; Lei et al., 2012; Li et al., 2014; Morris et al., 2013; Roberson et al., 2007; van Hummel et al., 2016). In contextual fear conditioning, both Fyn KO and DKO mice already showed deficits on training day, where the mice differed from WT at the 5^th^ (DKO) or 6^th^ (Fyn KO) minute (Fig. 2B). Thus, it is possible that Fyn depletion could have changed the way animals perceived pain, leading to a decrease in the saliency of the shock and weakening the association between shock and the chamber, with learning and memory being normal. Another possibility is that the expression of freezing behavior itself was impaired. Future studies using cue induced freezing or predator odor induced freezing would determine whether Fyn and/or tau depletion affected freezing behavior. In Fyn KO and DKO mice, the cognitive deficits were accompanied by decreases in Y1472-NR2B phosphorylation levels in the hippocampal insoluble PSD fractions while WT and tau KO mice had no memory deficits and no decreases in pY1472-NR2B. The similarity in the levels of pY1472-NR2B in our WT and tau KO mice agree with other findings using the same tau KO line used here (Lopes et al., 2016b). However, another report using a different tau KO line found reduced levels of pY1472-NR2B (Ittner et al., 2010). The fact that different lines of tau KO mice exhibited different phenotypes underlines the complexity of the analysis of tau KO mice (Ke et al., 2012; Lei et al., 2014).

In the brain, tyrosine phosphorylation of NR2B is important in regulating receptor trafficking, gating kinetics, and Ca^2+^ permeability (reviewed by (Groveman et al., 2012; Kalia et al., 2004; Trepanier et al., 2012)). Since Fyn KO and DKO mice had reduced levels of pY1472-NR2B, we examined calcium influx using hippocampal neurons of each genotype. As expected, Fyn KO neurons had reduced glutamate-induced Ca^2+^ response relative to WT neurons. Interestingly, DKO neurons had an even greater reduction relative to Fyn KO neurons while tau KO neurons had a response similar to WT neurons, suggesting that tau had a Fyn-independent role in regulating Ca^2+^ response that was unmasked in the absence of Fyn. Possible mechanisms include interactions of tau with other Ca^2+^ permeable channels such as voltage gated Ca^2+^ channels, regulators of NMDARs, or other components of the PSD, with tau modulating signaling pathways that increase intracellular Ca^2+^ levels. Additional studies are needed to explore these nonexclusive possibilities. Our Ca^2+^ response data from the tau KO neurons agree with a report showing that brain slices from tau KO mice had no change in glutamate-induced Ca^2+^ response profiles (Bi et al., 2017) and is consistent with the overall picture of the tau KO having no changes in excitatory post-synaptic currents (EPSCs), AMPA/NMDA current ratios, and LTP (Ittner et al., 2010; Kimura et al., 2014; Roberson et al., 2011). However, when tau was acutely depleted by shRNA in WT neurons, a decrease in Ca^2+^ response after glutamate stimulation was revealed (Miyamoto et al., 2017). This suggested that extended tau depletion, achieved through genetic ablation in a tau KO mouse, may be fundamentally different from acute tau depletion achieved through the use of tau shRNA. Indeed, although no permanent deficits in axonal development were reported in neurons from tau KO mice (Dawson et al., 2001; Harada et al., 1994), WT neurons treated with tau antisense oligonucleotides or shRNA exhibited dramatic deficits in axon formation defective migration properties, abnormalities in mitochondria transport, and developmental deficits (Caceres and Kosik, 1990); (Sapir et al., 2012). These findings indicate that comparing neurons from a knockout animal to those created by treating WT neurons with shRNA may be less straightforward than expected. Successful creation of a KO animal may be required to select genetic alterations that allow pups to be born and animals to be bred. Moreover, the extent of compensation in the tau KO mouse has been analyzed by microarray analysis and it was found that in the brain, several signal transduction genes such as fosB and c-fos were up-regulated 5-24 fold (Oyama et al., 2004). Based on these findings, we demonstrated that tau was required for NGF-induced AP-1 activation (Leugers and Lee, 2010). The fact that the loss of tau required up-regulation of both fosB and c-fos for compensation suggested that AP-1 activation was a critical tau function required by a mouse to survive.

Our finding that tau KO had no memory deficits, no changes in pY1472-NR2B, and no changes in glutamate induced Ca^2+^ response suggested that either Fyn did not require tau to mediate these functions or that another protein was compensating for the loss of tau with respect to its Fyn interaction. Interestingly, other microtubule-associated proteins (MAPs), MAP1A/1B and MAP2, are upregulated in tau KO mice (Harada et al., 1994; Ma et al., 2014) and in agreement with previous data reported for our tau KO mouse line (Ma et al., 2014), we found MAP2 to be increased in our tau KO mice (Fig. 6A). In contrast, MAP2 was significantly decreased in Fyn KO mice, correlating with the reduction and shortening of MAP2 positive dendrites described previously (Kojima et al., 1997). In DKO mice, MAP2 was decreased, indicating that the Fyn effect superseded the tau effect on MAP2 levels. The mechanisms by which Fyn depletion down-regulated or tau depletion up-regulated MAP2 are not known. However, since Fyn has been shown to affect protein translation (White et al., 2008) and tau, fos activation (Leugers and Lee, 2010), the effects of Fyn may be downstream of those of tau. Therefore, in the DKO, the Fyn KO trait resulted in the loss of MAP2-mediated compensation for the loss of tau whereas in the tau KO, up-regulated MAP2 levels would be capable of compensating for the loss of tau. In examining the glutamate induced Ca^2+^ response, the response in tau KO resembled those in WT due to compensation by MAP2. In contrast, with the reduction of MAP2 in both the Fyn KO and DKO and the attenuation of Ca^2+^ influx in the DKO relative to the Fyn KO, we were able to uncover a role for tau in the Ca^2+^ response that was independent of Fyn (Fig. 5C).

In the tau KO mice, if the tau-Fyn interaction was of critical importance, one would predict that compensation for the loss of the tau-Fyn interaction would occur. Our finding that tau KO neurons had a significantly higher density of MAP2-Fyn complexes in the dendrites relative to WT neurons suggested that MAP2 would have an increased role in interacting with Fyn in the absence of dendritic tau. We also showed that similar to tau, MAP2 also potentiated Fyn activity (Fig. 6B, C), leading us to hypothesize that MAP2 would have an increased role in interacting with Fyn, affecting NR2B phosphorylation and NR function in the absence of dendritic tau. Future studies using gain-of-function and loss-of-function experiments would further test this hypothesis.

In summary, we generated a tau^−/−^/Fyn^−/−^ DKO mouse and found that tau was capable of regulating Ca^2+^ influx in a Fyn-independent manner, a property that we propose to be mimicked by MAP2 in the absence of tau. As dysregulation of Ca^2+^ influx is a known mechanism leading to excitotoxity in AD, our findings identify tau as a new therapeutic target in the regulation of neuronal Ca^2+^. Most importantly, increased levels of dendritic MAP2-Fyn complexes in the tau KO highlight the importance of the tau-Fyn interaction and indicate the need to compensate if tau is lost. Our findings suggest that the association and activation of Fyn by tau in dendrites are critical neuronal functions.

## Methods and Materials

### Mice

Tau KO (C57BL/6) mice(Dawson et al., 2001) and Fyn KO (C57BL/6/S129) mice(Stein et al., 1992) were obtained from Jackson Laboratories and crossed to generate heterozygotes. The heterozygotes were then bred to generate tau^−/−^/Fyn^−/−^ and tau^+/+^/Fyn^+/+^ mice (Fig. S1). Mice were genotyped as described by Jackson Laboratories. All procedures were approved by the University of Iowa Institutional Animal Care and Use Committee and in strict accordance with the NIH Guide for the Care and Use of Laboratory Mice.

When the homozygous Fyn KO trait was bred into a C57BL/6 background, the mice exhibited a 50% chance of developing non-obstructive hydrocephalus postpartum (Goto et al., 2008). The DKO mice also developed hydrocephalus. Magnetic resonance imaging (MRI) was used to differentiate normal from hydrocephalic mice. Only normal mice screened by MRI were used for behavioral and biochemical tests.

### Magnetic resonance imaging

Varian Unity/Inova 4.7 T small-bore MRI system (Varian, Inc., Palo Alto, CA; Small Animal Imaging Facility, University of Iowa) with an in-plane resolution of 0.13 × 0.25 mm^2^ and 0.6 mm slice thickness was used. Coronal images were collected and lateral ventricle volume sizes were analyzed with ImageJ (Carter et al., 2012). A lateral ventricle volume ≤0.2 units was normal, ≥0.2 and ≤ 2 units was moderate, and ≥ 2 units was severe hydrocephalic.

### Open field test

The open field test was done at 8 weeks as described(Coryell et al., 2007) with a dimension of 40.6 × 40.6 × 36.8 cm open-field chamber (San Diego Instruments, San Diego, CA), 55 lux, for 20 min. Total activity was measured by total beam breaks and central activity was measured by beam breaks in center (15.2 × 15.2 cm). 14 WT (14 male), 20 Tau KO (9 male, 11 female), 18 Fyn KO (15 male, 3 female), 13 (10 male, 3 female) DKO mice were tested. Since there were no statistically significant differences between males and females, their results were combined.

### Pole test

The pole test was done at 12 months of age as described(Lei et al., 2014). Trials were excluded if the mouse jumped or slid down the pole. Between tests, the pole was cleaned using 70% ethanol. 9 WT, 13 tau KO, 11 Fyn KO, 18 DKO male mice were tested.

### Novel object recognition

The novel object recognition test was done at 10 weeks as described(Tang et al., 2001). The time each animal spent exploring objects (defined as nose within an inch of the object) was manually determined. Discrimination index [DI = (X_1_-X_2_) / (X_1_+X_2_)] was calculated. 17 WT (12 male, 5 female), 17 tau KO (12 male, 5 female), 28 Fyn KO (16 male, 12 female), 18 DKO (14 male, 4 female) mice were used. Since there were no statistically significant differences between males and females, their results were combined.

### Contextual fear conditioning

Contextual fear conditioning was done at 11 weeks as described(Sowers et al., 2013), using a near-infrared video-equipped fear conditioning chamber (Med Associates, Inc., St. Albums, VT). Freezing was scored with VideoFreeze software (Med Associates, Inc.). 32 WT (27 male, 5 female), 29 tau KO (21 male, 8 female), 21 Fyn KO (14 male, 7 female), 20 DKO (14 male, 6 female) mice were tested. Since there were no statistically significant differences between males and females, their results were combined.

### Pentylenetetrazole-induced seizures

Pentylenetetrazole (PTZ)-induced seizure was done at 12 weeks as described(Roberson et al., 2007). A modified Racine seizure scale was used: 0: normal behavior; 1: immobility; 2: tail extension; 3: forelimb clonus; 4: generalized clonic activity; 5: bouncing; 6: tonic extension; 7: death(Loscher et al., 1991; Racine, 1972). 25 WT (14 male, 11 female), 21 tau KO (9 male, 12 female), 36 (25 male, 11 female) Fyn KO, and 26 DKO (19 male, 7 female) mice were used. Since there were no statistically significant differences between males and females, their results were combined.

### Tissue preparation and subcellular fractionation

PSD preps were isolated from hippocampi of 9-12 months old mice(Ittner et al., 2010; Lopes et al., 2016b; Milnerwood et al., 2010). As described, crude synaptosomal membranes were resuspended with 1% Triton X-100 containing buffer and after centrifugation, the supernatant containing non-PSD membranes was retained as the “soluble non-PSD” fraction. The pellet was resuspended in 1% Triton X-100, 1% deoxycholic acid, 1% SDS containing buffer and after centrifugation, the supernatant was retained as the triton “insoluble PSD” fraction.

### Primary hippocampal neuron culture

Primary hippocampal neuronal culture was performed as described(Beaudoin et al., 2012). Hippocampi from P0 pups were isolated and digested in 1 mg/ml trypsin (Sigma-T7409) and sequentially triturated with Pasteur pipettes. Cells were plated with MEM media (Gibco) containing 10% horse serum and 1µM insulin (Sigma-I5500) and maintained with Neurobasal-A Plus and 2% B27 Plus (Gibco) at 37 °C and 5% CO_2_ until harvest. 50% of media was changed every 3 days.

### In situ proximity ligation assay (PLA) and immunofluorescence

PLA components were purchased from Sigma (Duolink^®^ *in situ*) and the assay was performed per manufacturer’s instructions. Cells were imaged using confocal microscopy (Leica SP8 STED) and exported to ImageJ for analysis.

Primary antibodies used were: DA9 (1:50, mAb, generous gift of Dr. Peter Davies), Fyn3 (sc-16, 1:250, Santa Cruz), MAP2 (1:1000, chicken polyclonal, Thermo Fisher, Cat **#** PA1-10005), MAP2 (HM-2, 1:250, Sigma M4403), Src (36D10, 1:250, rabbit polyclonal, Cell Signaling, Cat #2109), phospho-Src (1:250, rabbit polyclonal, Tyr418; Thermo Fisher, Cat #44660G), and tubulin (1:50, YL1/2; Accu-Spec). Secondary antibodies used were anti-mouse Alexa 488 (1:250, Molecular Probes), anti-chicken Alexa 633 (1:1000, Thermo Fisher), and anti-rat Alexa 647 (1:250 Jackson Immunoresearch).

To quantify the density of PLA puncta, ten random areas on each coverslip were photographed. Axons were identified by either tau or tubulin positive staining and MAP2 negative staining; dendrites were identified by MAP2 positive staining. The lengths of all visible processes were measured using ImageJ and all visible PLA puncta were manually counted. For each of the 10 areas, the densities of dendritic or axonal PLA puncta were calculated by dividing the number of PLA puncta located on dendrites or axons by the total dendrite or axon length, respectively. The experiment was performed 3 times. In Fig. 3, the density of PLA in dendrites was expressed relative to the density in axons, where the average value was set as “1”. In Fig. 7, the density of PLA in tau KO mice was expressed relative to the density in WT mice, where the average value was set as “1”. In Fig. S3, for each neuron, the proximal axon length was defined as the average length of the dendrites from that neuron.

### Ca^2+^ imaging

Ca^2+^ imaging was performed on primary hippocampal neurons (DIV 11-12) as described(Kim et al., 2009). Hippocampal neurons were loaded with 2µM Fura-2/AM (Invitrogen) and then mounted on an inverted microscope (Olympus IX-71). The cells were perfused with HEPES buffered Hank’s salt solution (HH buffer) to establish baseline and then stimulated with 100 µM Glutamate, 10 µM Glycine, and 0.2 µM Tetrodotoxin (TTX) in HH buffer. [Ca^2+^]_*i*_ changes were continuously recorded by alternately exciting fluorescence at 340 nm (12-nm bandpass) and 380 nm (12-nm bandpass) using a Polychrome V monochromator (TILL Photonics, Munich, Germany). Fluorescence was recorded at 530 nm (50-nm bandpass) using an IMAGO charge-coupled device camera (640 × 480 pixels; TILL Photonics). A 20x objective (numerical aperture = 0.75, Olympus, Japan) and a 2×2 binning set at room temperature was used. [Ca^2+^]_i_ was derived using the formula [Ca^2+^]_i_ = *K*_*d*_β(*R* − *R*_min_)/(*R*_max_ − *R*), with *K*_*d*_=275nM (Shuttleworth and Thompson, 1991), *R* =*F*_340nm_/*F*_380nm_, *R*_min_= 0.21, *R*_max_= 3.45, and β= 6.97. Data were analyzed using TILLvisION version 4.0.12 software (TILL Photonics).

### Immunoprecipitation

3T3 (NIH) cells were grown in α-MEM supplemented with 10% fetal bovine serum (Hyclone) and transfected with either Fyn alone or with Fyn and MAP2c together. Cells were harvested with RIPA lysis buffer. Fyn was IPed with 1µg Fyn antibody (mouse monoclonal antibody [mAb], sc-434, SantaCruz) and 10 µL Protein G plus-agarose (CalBiochem). Washed Protein G bead pellets were resuspended in 2X Laemmli buffer and subjected to Western blot analysis. 3T3 cells have not been reported as contaminated or misidentified (International Cell Line Authentication Committee, October 14, 2018).

### Plasmids

pRc/CMV expressing the N-terminal fragment of human MAP-2c was synthesized by site-directed mutagenesis using the pRc/CMV MAP2c plasmid (Albala et al., 1993) as template. Residue 320 on MAP-2c was mutated to a stop codon. The construct was sequenced to verify the MAP2c sequence and the mutation site.

### *In vitro* phosphorylation of tubulin

*In vitro* phosphorylation of tubulin was performed as described (Sharma et al., 2007). Fyn or truncated MAP2c (human N-MAP2c residue 1-319) were expressed in 3T3 cells and IPed using 1µg anti-Fyn (mAb, sc-434, SantaCruz) or 4 µg anti-MAP2 (HM-2, mAb, Sigma, Cat # M4403) and Protein G beads (CalBiochem). The beads containing Fyn were resuspended in 100 µL buffer (0.1% Triton, 50mM Tris pH7.5, 150 mM NaCl, 1 mM EDTA, 1mM AEBSF) and equally divided into two tubes. N-MAP2c beads were added to one of these tubes and nutated at 4°C for 1 hour, centrifuged, and supernatant discarded. Control tube was similarly treated, where Protein G beads were added to beads containing Fyn. 10 µg taxol-stabilized tubulin and kinase reaction buffer was added to both tubes and incubated for 5 min at 37 °C. A third tube containing only tubulin, kinase reaction buffer, and 1µg mouse non-specific IgG was used as control. Samples were subjected to western blot analysis.

### Western blot analysis

Western blotting was performed as described(Lee et al., 1998). Primary antibodies used were: Tau5-HRP (1:10,000, (Leugers and Lee, 2010)); Fyn3 (sc-16, 1:1000, rabbit polyclonal, Santa Cruz); GAPDH (1:25,000, mAb, Chemicon, Cat # MAB374); synaptophysin (SP15, 1:1000, mAb, Millipore, Cat # MAB329-C); PSD-95 (EP2652Y, 1:1000, rabbit monoclonal, Millipore, Cat# 04-1066); NR2B (1:1000, mAb, One World Labs, StressMarq, Cat # SMC-33D); phospho-Y1472-NR2B (1:1000, rabbit polyclonal, PhosphoSolutions, Cat # p1516-1472); Src (GD-11, 1:1000, mAb, Milipore, Cat # 05-184); phospho-SFK (1:1000, phospho-Src Tyr 418, rabbit polyclonal, Cell Signaling, Cat # 2101); MAP2 (HM-2, mAb, 1:1000, Sigma, Cat # M4403); generic phospho-tyrosine (4G10, 1:1000, mAb, Millipore, Cat # 05-321); and tubulin (1:1000, YL1/2; Accu-Spec). Quantification was done by densitometry using ImageJ. Protein levels from the PSD fraction were normalized to β-actin (Fig. 4) and protein levels from crude lysates were normalized to GAPDH (Fig. 6A), with values subsequently normalized to WT levels, which were represented as “1”. In Fig. 6C, pSFK was normalized to total Fyn and values from Fyn+MAP2c were normalized to Fyn alone condition with the latter represented as “1”. In Fig. 6D, 4G10 was normalized to both tubulin and Fyn and the Fyn+N-MAP2c condition was normalized to the Fyn alone condition with the latter represented as “1”.

### Statistical Analysis

Statistical analysis was carried out with either Microsoft Excel or GraphPad Prism 7.0 using ordinary one way ANOVA with Tukey’s post hoc multiple comparison or unpaired parametric t-test when appropriate. For Fig. 2B and 2E, 2-way ANOVA with Tukey’s post hoc multiple comparison was used. p≤0.05 was considered as statistically significant, with p≤0.05 denoted as *, p≤0.01 denoted as **, p≤0.001 denoted as ***, and p<0.0001 denoted as ****.

## Declarations

### Animals

Animal care was in compliance with National Institutes of Health guidelines for the care and use of laboratory animals and all animal procedures were approved by the University of Iowa Institutional Animal Care and Use Committee.

### Availability of data and material

The datasets analysed in the current study are available from the corresponding author on reasonable request.

### Authors’ contributions

G Liu designed the experiments, performed the PCR, behavioral, biochemical, calcium imaging, and PLA experiments, analyzed data, and wrote the manuscript. R Thangavel and E Adams performed IHC and analysis, MB Francis generated the double knockout mice, Y Kim assisted with PLA experiments, J Rysted and Z Lin assisted with calcium imaging, RJ Taugher assisted with behavioral work, and JA Wemmie and YM Usachev provided valuable discussions and data analysis. G Lee designed experiments, analyzed data, and revised the manuscript.

## Acknowledgements

We thank Drs. Peter Davies for generously providing the DA9 antibody, Stefan Strack for valuable suggestions, Bridget Shafit-Zagardo for MAP2c plasmid, Dan Thedens for help with MRI, Skye Souter for technical assistance, and Craig Morita for help with figures. We acknowledge the University of Iowa Medical Scientist Training Program for support to Guanghao Liu and the Holden Comprehensive Cancer Center (NIH/NCI P30CA086862) for support at the University of Iowa Central Microscopy Research Facility and Small Animal Imaging Facility.

